# Sperm-particle interactions and their prospects for charge mapping

**DOI:** 10.1101/624510

**Authors:** Veronika Magdanz, Johannes Gebauer, Priyanka Sharan, Samar Eltoukhy, Dagmar Voigt, Juliane Simmchen

## Abstract

In this article, we demonstrate a procedure to investigate sperm charge distribution by electrostatic sperm-particle interactions. We fabricated and investigated differently charged particles and their attachment distribution on the bovine sperm membrane. We observed the sperm-particle attachment sites using bright field and cryo-scanning electron microscopy combined with energy-dispersive X-ray analysis. Our findings suggest that the charge distribution of the sperm membrane is not uniform and although the overall net charge of the sperm cell is negative, positively charged areas are especially found on the sperm heads. We test the newly developed method to investigate the dynamic charge distribution of the sperm cell membrane upon maturation induced by heparin, as a representation of the multitude of changes during the development of a sperm.

---

Colloidal chemistry has developed a wide variety of techniques allowing the synthesis of particles and the modification of their physicochemical properties (material, size, charge, shape, stiffness, hydrophobicity) for the desired cell-particle interactions.^[1]^ Numerous biomedical science areas have benefited from the interaction between cells and micro- and nanoparticles. This has paved the way for applications of cell-particle interactions in nanomedicine, for cell separation, imaging and actuation.^[1–6]^ Additionally, cell-particle interactions are increasingly used to study cell mechanics and physiology as well as immune response.^[3]^

Likewise, in reproductive technologies, nano- and microparticles have been proven useful for the enrichment of high-quality sperm by MACS (magnetic-assisted cell sorting) and imaging and tracking of sperm in vitro and in vivo.^[7,8]^ Additionally, nanoparticles can mediate the drug and gene delivery into sperm.^[9]^ Recently, the loading of iron oxide nanoparticles, besides other payloads, into sea squirt sperm cells has led to the development of chemotactic sperm-driven micromotors.^[10]^

In these existing techniques, little attention has been paid towards the electrostatic interaction between the particles and sperm cells. The binding between particles and sperm cells has been achieved mainly by specific cell receptors such as antibodies, or cell internalization agents.^[7]^ Sperm cells, as most mammalian cells have a net negative surface charge (NNC).^[11]^ Microelectrophoresis has been used to isolate sperm with a NNC for assisted reproduction techniques, which were found to be mature cells (not capacitated) and more likely to contain intact DNA compared to sperm with a net positive or neutral charge.^[12,13]^ The interaction of sperm cells with their surroundings plays a major role in their journey to the fertilization site. It is known that sperm cells interact with epithelial cells in the oviduct and that molecular interactions with the female reproductive tract allow the passage through. During their development, sperm cells alter their membrane charge by interaction with the surrounding fluids. During the epididymal maturation, sialic acid, sialoglycoproteins, steroid sulfates and sulfated carbohydrates are incorporated into the sperm surface, where they act as stabilizers for the membrane and increase the NNC.^[14]^

Upon maturation of the sperm, a decrease of the NNC occurs, accompanied by increasing the permeability of the membrane to calcium, which promotes acrosome reaction.^[14]^ A decline of the NNC of sperm is also observed by incubation with follicular fluid or neuraminidase, which proves the interaction between the sperm cell and the surrounding medium. A reduction of the NNC seems to be required for the sperm passage through the cumulus cell matrix and binding to the zona pellucida. This illustrates that the sperm cell charge is a dynamic property, which changes during the development of the sperm cell. Also, due to the characteristic morphology of the sperm with distinct functions of certain parts of the cell (acrosome – fusion with egg, head – contains DNA, midpiece – mitochondrial sheath, tail – motility), the surface charge is not uniform across the membrane. The abrupt differences between morphologically distinct segments suggest that the biochemical properties of the plasma membrane also vary between these areas.^[11]^ More than most other cell types, the properties of the sperm membrane are very important regarding all functions and interactions of the male gamete, e.g. motility, sperm survival in the male and female reproductive tract, interaction with the epithelium of the oviduct, oocyte fusion, et cetera.^[15]^ Thus, it is suspected that a charge map of sperm cells will give insight on their functional integrity and state and allow inference regarding their migration and fertilization capability.

Here, we investigate the interaction of bovine spermatozoa with a range of micro- and nanoparticles. The particles differ in surface charge (quantified by zeta potential) and are composed of different materials (silicon dioxide, ferric oxide or both) (Table 1). The attachment of particles on the membrane of the sperm cells is studied in dependence on their surface charge. The distribution of the particles across the sperm membrane is investigated in order to explore their potential for creating a charge map of the sperm cell. The proof-of-concept of charge mapping of sperm is demonstrated by brightfield microscopy of sperm binding to differently charged particles and by the combination of cryo-SEM and EDX (cryo-scanning electron microscopy and energy-dispersive x-ray analysis) of sperm binding to particles consisting of different elements.

**Table 1.** Particles, their shape, range of dimensions and surface charges (positively charged: orange; negatively charged: blue). H1-H4 are hematite particles. Surface charge in H_2_O was measured by zetasizer (dynamic light scattering, see Figure S1).

| Type of particles | Core material | Coating | Shape | Size [nm] | Surface charge in $\text{H}_2\text{O}$ [mV] |
| --- | --- | --- | --- | --- | --- |
| H1 | $\text{Fe}_2\text{O}_3$ | - | spherical | 1000 | 46.1 |
| H2 | $\text{Fe}_2\text{O}_3$ | - | spherical | 5 - 10 | 12.9 |
| H3 | $\text{Fe}_2\text{O}_3$ | - | spherical | 100 | 14.8 |
| H4 | $\text{Fe}_2\text{O}_3$ | | elongated | 100 | 12.9 |
| $\text{SiO}_2$@H1 | $\text{Fe}_2\text{O}_3$ | Silica | spherical | 1000 | -27.8 |
| $\text{SiO}_2$@H4 | $\text{Fe}_2\text{O}_3$ | Silica | elongated | 100 | -18 |
| $\text{SiO}_2$ | Silica | - | spherical | 1000 | -39.5 |

We selected different particles having a range of zeta potential in the positive and negative range (see Table 1). The particles cover a size range from 5 nm to 1 μm. The particles are either composed of iron oxide (H1-H4, positively charged) or silica (SiO_2_, negatively charged) or iron oxide particles covered with silica (SiO_2_@H1, SiO_2_@H4, negatively charged).

The attachment of particles to the bovine sperm cells was observed under bright field microscope after incubating the sperm cells with the particles for 30 minutes. Details on the protocol can be found in the experimental section. The experiments were repeated 3-5 times, and each time more than 20 images were randomly taken from which over 100 sperm-particle constructs were analyzed. Additionally, cryo-SEM was performed. For the preparation of the samples, the sperm cells were washed twice in water and mixed with the microparticles. A droplet of 1 μL volume of this suspension was pipetted onto a clean glass cover slide fixed to the cryo-SEM sample holder, shock-frozen in liquid nitrogen, transferred to the −140 °C-cold cryo preparation chamber, sublimed for 10-15 minutes at −70 °C, and the protocol was further followed as described previously by Bräuer et al.^[16]^.

Figure 1 displays representative images of positively charged particles bound to bull sperm. It is visible that particles of all dimensions bound to the sperm surface, while the smaller particles (Figure 1i & ii) tended to cover the whole area of the membrane. The elongated particles (Figure 1iii) showed tendency to agglomerate, which is due to the parallel alignment of magnetic moments of these iron oxide particles. Our measurements of the bovine sperm surface zeta potential by two different techniques (dynamic light scattering and microelectrophoresis) revealed that the surface zeta potential of bovine sperm lies in the range of −10 to −30 mV (Experimental section and Figure S1 in Supporting information). Thus, electrostatic interactions between the bull sperm membrane and positively charged particles were favored.

**Figure 1:**
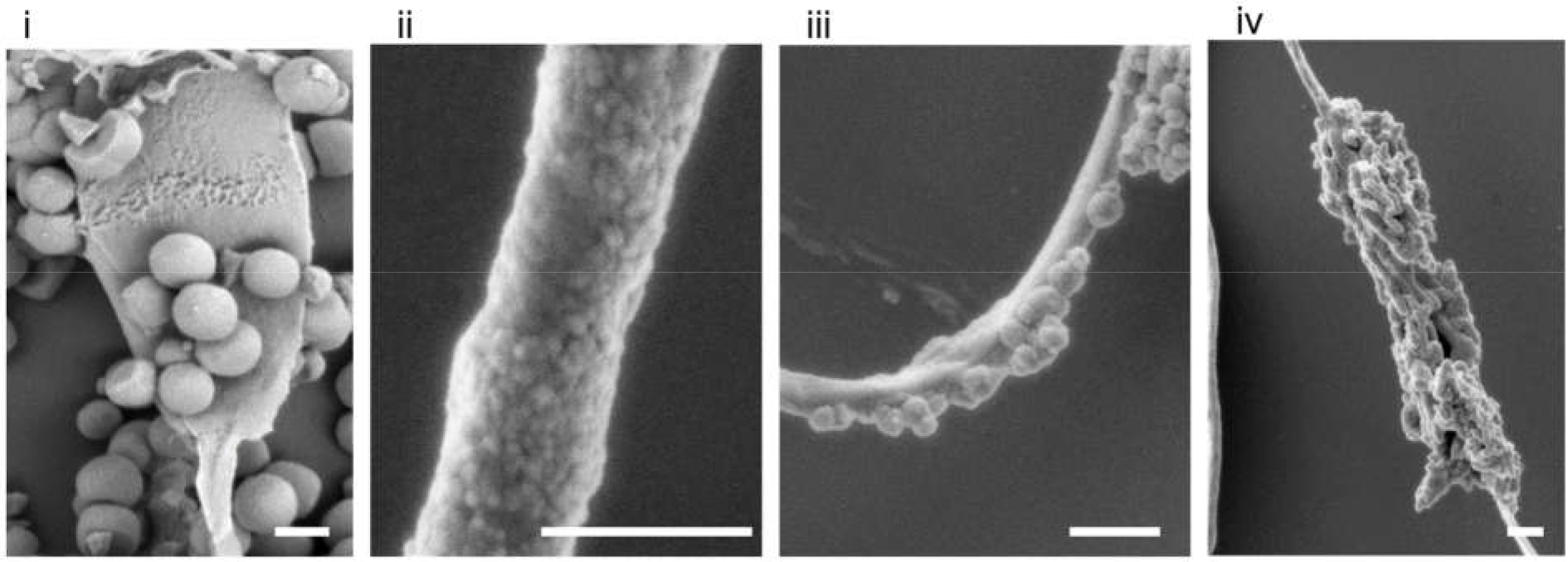
Cryo-SEM images showing positively charged micro/nanoparticles on the surface of bovine spermatozoa. Particle types: (i) H1, (ii) H2, (iii) H3, (iv) H4. Scale bars are 1 μm.

Surprisingly, negatively charged particles (Figure 2) also bound to sperm cells on some areas of their membrane. This hints towards the presence of positively charged areas on the sperm surface. Even though the overall surface charge of sperm cells is negative, it is possible that some areas on the sperm’s membrane display a positive charge, due to the loss of integrated negatively charged glycoproteins.^[12]^ Additionally, the fusion with the oocyte has been described as a membrane potential-depending event, in which the egg performs a quick change from negative to a positive potential as an electrical block against polyspermy^[17,18]^ (entering of a second sperm after fusion).

**Figure 2:**
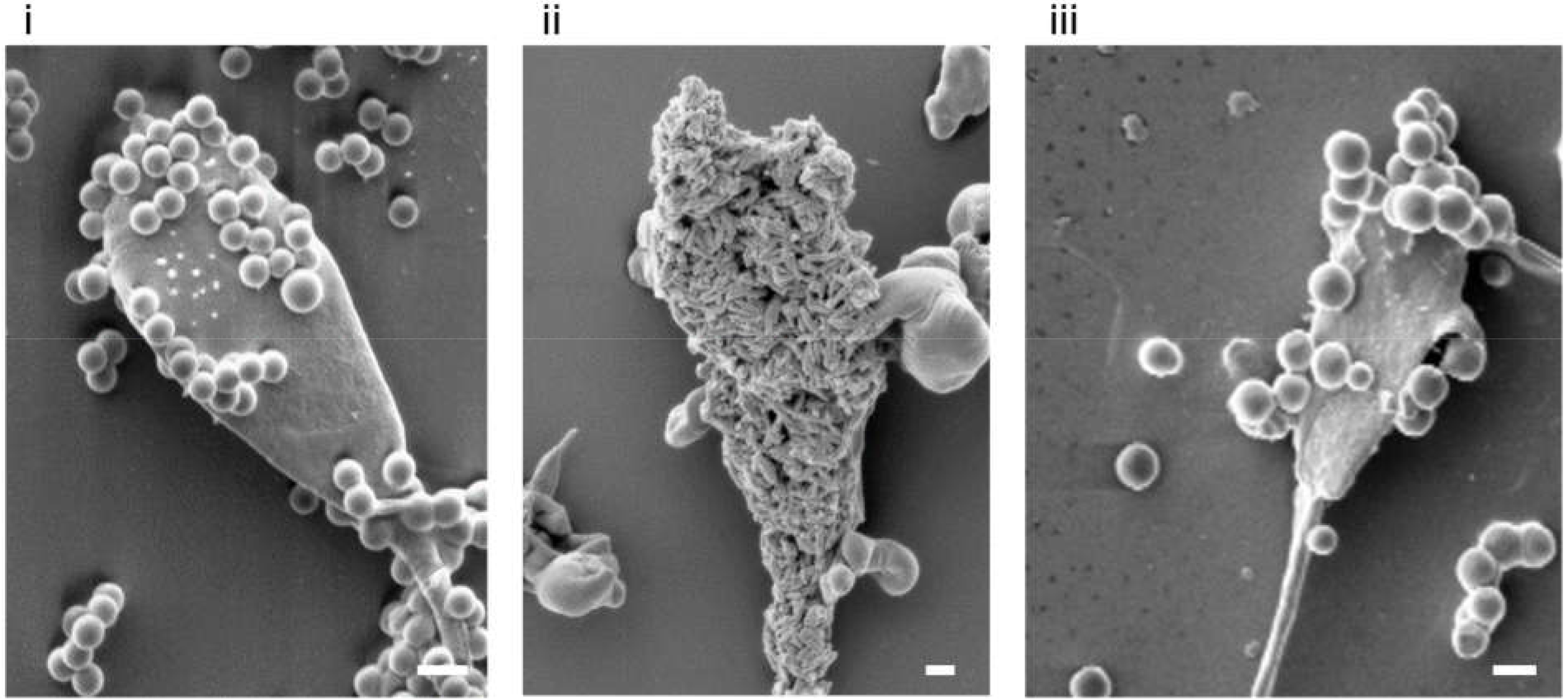
Cryo-SEM images showing negatively charged micro/nanoparticles attached to the surface of bovine spermatozoa. Particle types: (i) SiO_2_@H1 (ii)SiO_2_@H4, (iii) SiO_2_. Scale bars are 1 μm.

Figure 3 gives a more detailed overview of the distribution of the attached particles. It is very apparent that negatively charged particles bound more frequently on the sperm heads (red bars in Figure 3ii) and only rarely on the flagellum or the whole cell (this was found in less than 7% of the cases). Positively charged particles bound only to sperm heads in about 32% of the cases (Figure 3i), thus significantly less than negatively charged particles (52%, p=0.000002, oneway ANOVA, Bonferroni multiple comparison method). The attachment of positively charged particles to the flagella of the sperm cells occurred on average in 13% of the cases and more likely than negatively charged particles (6%, p=0.037). The attachment onto the whole body of the cell occurred slightly more frequently with positively charged particles (Figure 3i, yellow bars), but not significantly (p=0.09).

**Figure 3:**
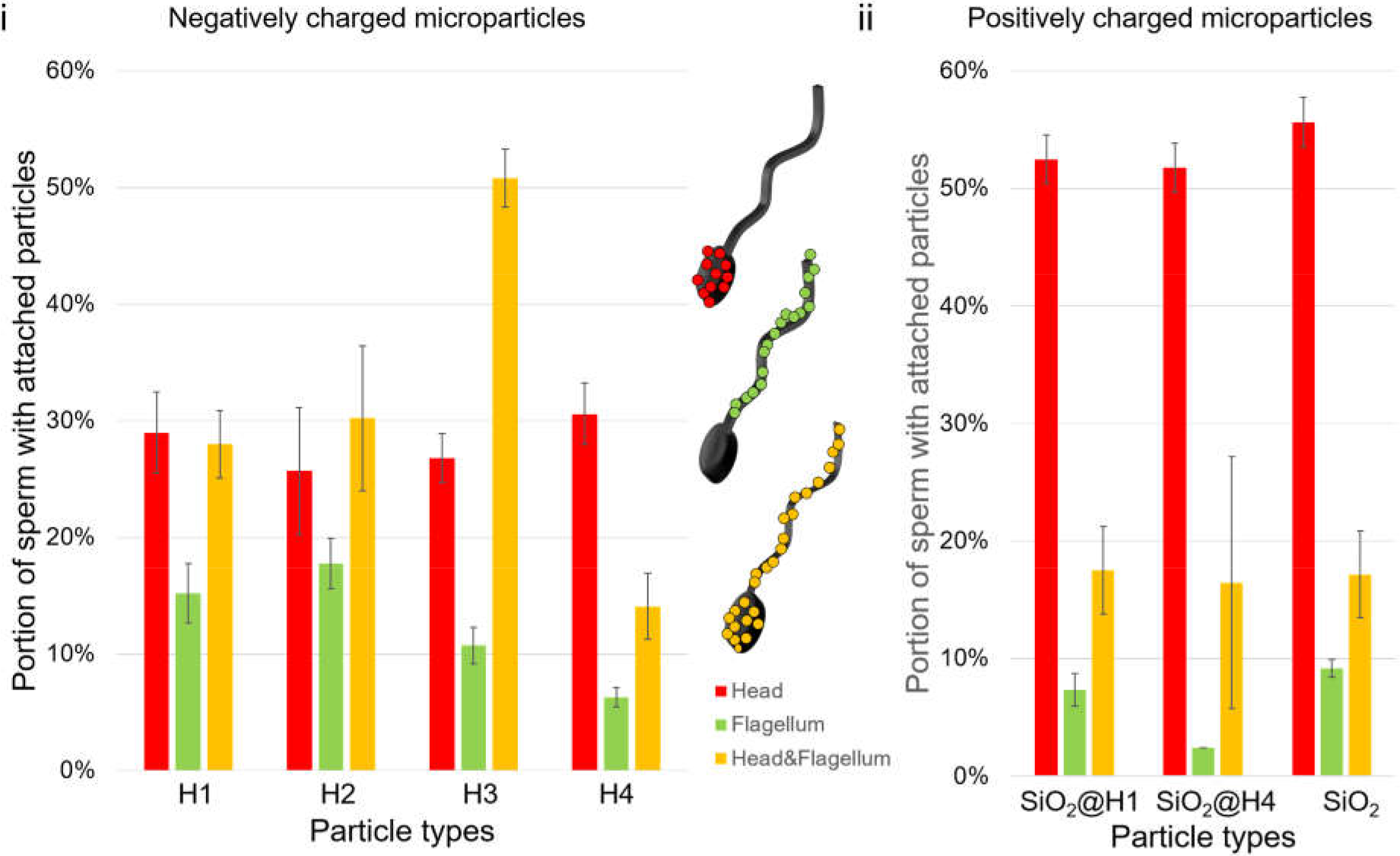
The graphs illustrate the distribution of positively charged (i) and negatively charged (ii) particles attached to the head (red), flagellum (green) or head & flagellum (yellow) of the spermatozoa. Columns correspond to means and bars to standard error of the mean; n= 3-5 measurement runs with >290 cells each.

These data suggest that bovine sperm contain a positively charged area on their heads which allowed the electrostatic binding with negatively charged particles. This rarely occurred on the sperm tails, where the chance of binding negatively charged particles is below 10%.

We performed sperm viability tests by staining of the bovine sperm cells with the sperm viability kit (Thermo Fisher) including SYBR14 (green) and propidium iodide (red), see Figure 4i+ii in order to evaluate the toxicity of the particles to the sperm.

**Figure 4:**
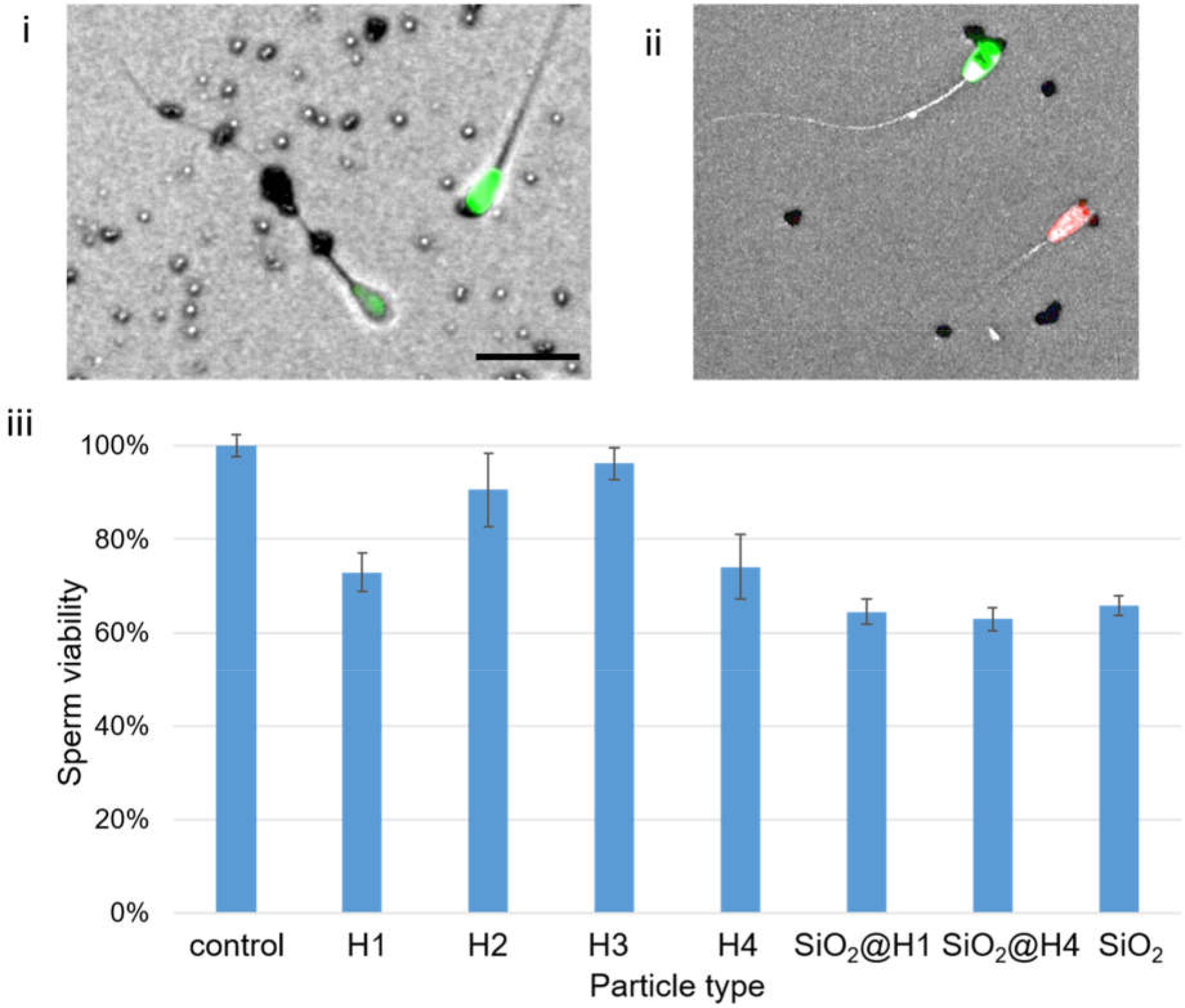
Cell viability of bull sperm in contact with microparticles tested with Live/dead viability stain. Green (SYBR14) stains live cells, red (propidium iodide) stains dead cells. Exemplary overlayed images (bright field, red and green channel) for particle types i) H1 and ii) SiO2@H1. Scale bars 20 μm. The bar chart in iii) compares the percentage of live sperm in all microparticle-sperm suspensions. Values for particle-covered sperms are normalized to the control. Columns correspond to means and bars to standard error of the mean. A total N>20,000 cells were analyzed with n >530 cells for each bar.

The graph in Figure 4iii displays the viability of sperm bound to particles normalized to the viability without particles (control). This normalization was performed, because thawed cryo-preserved semen samples always contain a certain amount of dead cells and we only wanted to consider the influence of the particle binding on the viability. As illustrated in Figure 4, the viability was maintained to over 60% of the initial sperm viability for all types of particles. The reduction of viability upon addition of microparticles was not significant (one-way ANOVA, Bonferroni multiple comparison method, p>0.05 for all particle types).

The cell viability test was also performed to allow a differentiation between the particle distribution patterns on live and dead sperm. The particle attachment on sperm heads showed no significant difference between live and dead sperm (Figure 5i). Live sperm cells seemed to bind significantly more frequently positively charged H1, H3 and H4 particles on their flagella (Figure 5ii, p=0.02, 0.001, 0.02, respectively) compared to dead sperm. Additionally, the negatively charged particles SiO_2_@H4 also bound more frequently on live sperm (p=0.046). Figure 5iii shows that dead sperm bound particles more often on their whole body surface compared to live sperm. This effect was significant for particle types H2 and H3 as well as SiO_2_@H1 (p=0.01, 0.03, 0.04, respectively).

**Figure 5:**
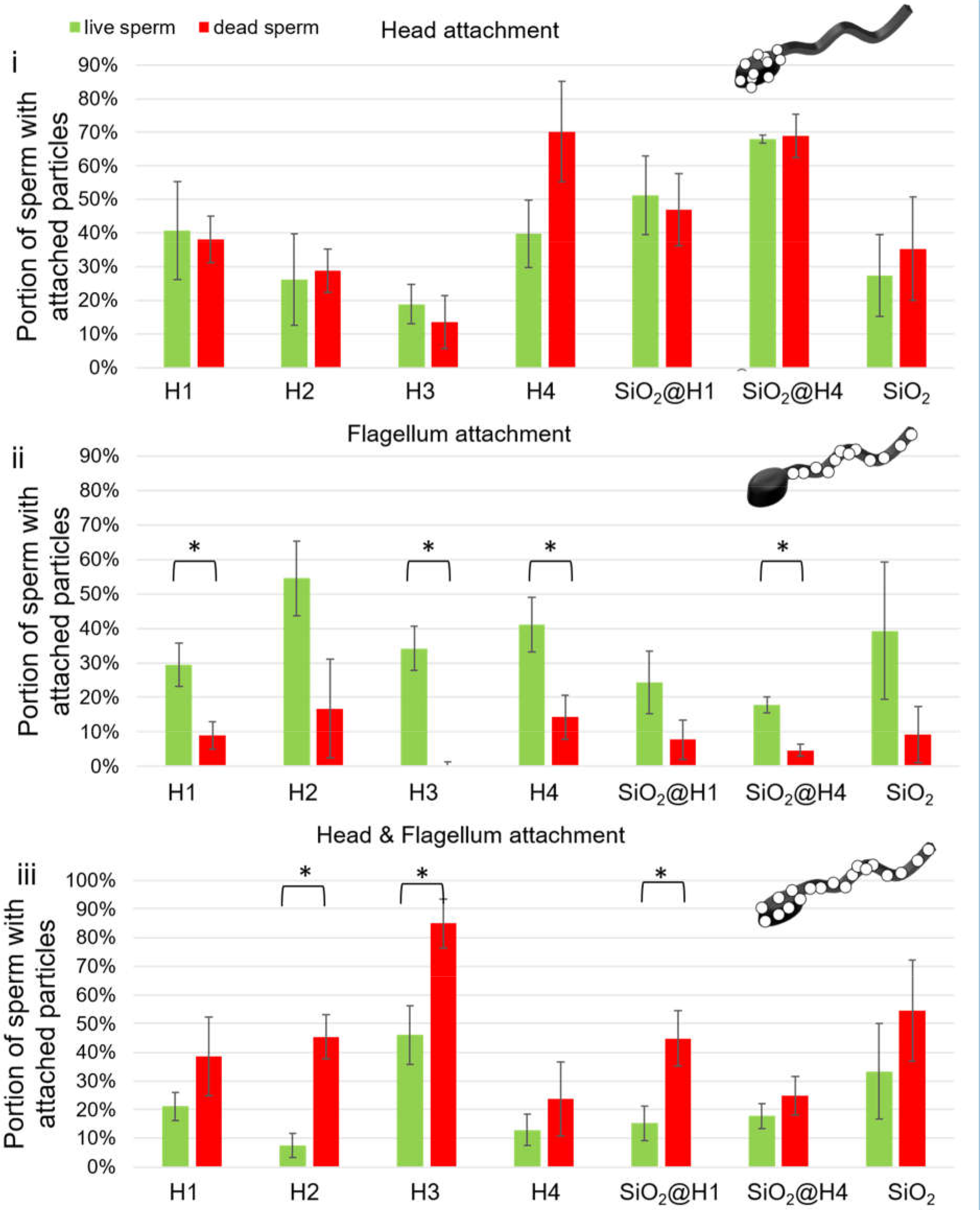
Comparison of the portions of sperm with attached particles divided between live and dead bull sperm cells on a) head, b) flagellum and c) whole cell body (head & flagellum). Columns correspond to means and bars to standard error of the means. A total number of 2850 cells were analyzed with at least 290 for each type of particle.

This could originate from a change in membrane potential upon apoptosis.^[19]^ It is not known how exactly the membrane charge changes during apoptosis, but it is expected that the NNC dissipates upon initiation of mitochondrial membrane permeabilization.^[20]^

Finally, we demonstrate the potential of sperm-particle interactions for studies of the charge distribution on sperm cells. By the incubation of sperm cells with two types of particles distinctly different in surface zeta potential, we investigated the particle distribution on the sperm’s surface by bright field microscopy. In order to prove that this method can distinguish changes in the membrane charge that occur throughout the sperm’s lifetime, we compared particle attachment on maturated and non-maturated cells. Maturation is an essential ripening process of spermatozoa prior to fertilization and includes the capacitation and hyperactivation of spermatozoa. It can be induced in vitro in bovine sperm cells by heparin ^[21,22]^. This maturation process is known to induce reorganization of lipids and proteins in the sperm’s membrane^[23]^.

A representative image of a non-maturated bovine sperm cell is displayed in Figure 6i, with silica particles mostly bound the head and a few iron oxide particles on head and flagellum. Figure 6ii displays a representative case of maturated sperm cell, which has significantly more iron oxide particles bound to its head and tail. In contrast, the negatively charged silica particles did not bind to the sperm’s head. Figure 6iii shows a fluorescent image of the same sperm cell as Figure 6ii with a fluorescent signal on the mid-piece and head of the sperm cells originating from the heparin-fluorescein conjugate that was used to mature the cell prior to particle binding. The portion of maturated and non-maturated sperm cells with attached particles was quantified and results are displayed in the bar graphs in Figure 6. As shown in the graph in Figure 6iv, the negatively bound silica particles preferentially bound on the sperm’s head, while the positively charged iron oxide particles were found on the whole body of the sperm cell. This was in agreement with the single-particle-type binding assays discussed above and displayed in Figure 3.

**Figure 6:**
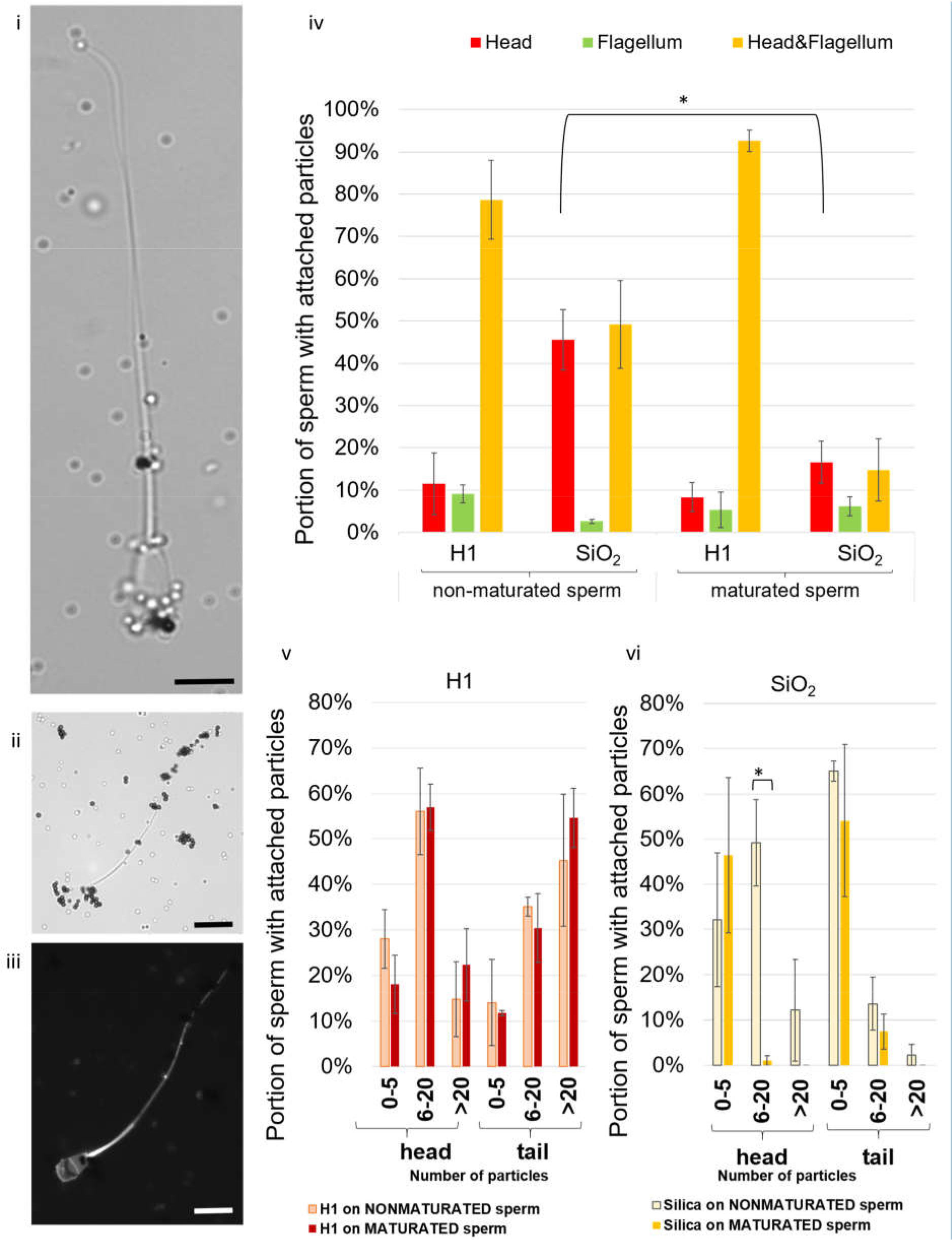
Brightfield microscopic images of iron oxide (black particles) and silica particles (bright particles) attached to i) nonmaturated bovine sperm and ii) maturated bovine sperm cells. iii) Fluorescent image of maturated sperm cells shows uptake of heparin (fluorescein conjugate) by head and midpiece of sperm. Scale bars 10 μm. iv) Bar graphs of portion of non-maturated and maturated sperm with attached iron oxide and silica particles. Positively charged H1 bind mostly on whole cell (>70%), while silica particles about 50% to head or whole cell on non-maturated sperm, versus >90% iron oxide and <20% silica particles binding to head and tail of maturated cells. Each bar is derived from three replicate measurements of at least 50 cells. One-way ANOVA followed by Bonferroni’s multiple comparison method reveal significant difference for the portions of maturated versus non-maturated sperm cells (p=0.03) attached to silica particles. v) shown by number of particles attached to head or tail of maturated and non-maturated sperm cells. In addition, statistical evaluation shows significant lower portion of silica particles binding to heads (p=0.007).

It is shown in Figure 6iv that a reduced silica particle binding occurred upon maturation of the sperm cells. When analyzing the number of attached silica particles (Figure 6vi), a drastic decrease of bound silica particles to the sperm head was clearly observed. Binding to the whole cell (head and flagellum) was reduced from 48% to 15%, but not significant (p=0.0055, Bonferroni’s multiple comparison method subsequent to one-way ANOVA). In contrast, the binding of positively charged iron oxide particles did not change significantly its pattern (Figure 6v).

In addition to the charge mapping with microparticles in brightfield microscopy, we also studied the cell-particle interactions with particle types H1 (positively charged iron oxide) and silica (negatively charged) using cryo-SEM and EDX. The EDX allows the identification of the two same-sized particles via element analysis. Figure 7 illustrates the results by representative images, which confirm the obtained results by brightfield microscopy (Figure 3). Silica particles (Figure 7: orange) attached preferably to the sperm heads, whereas the H1 (green) particles attached to the whole sperm body.

**Figure 7:**
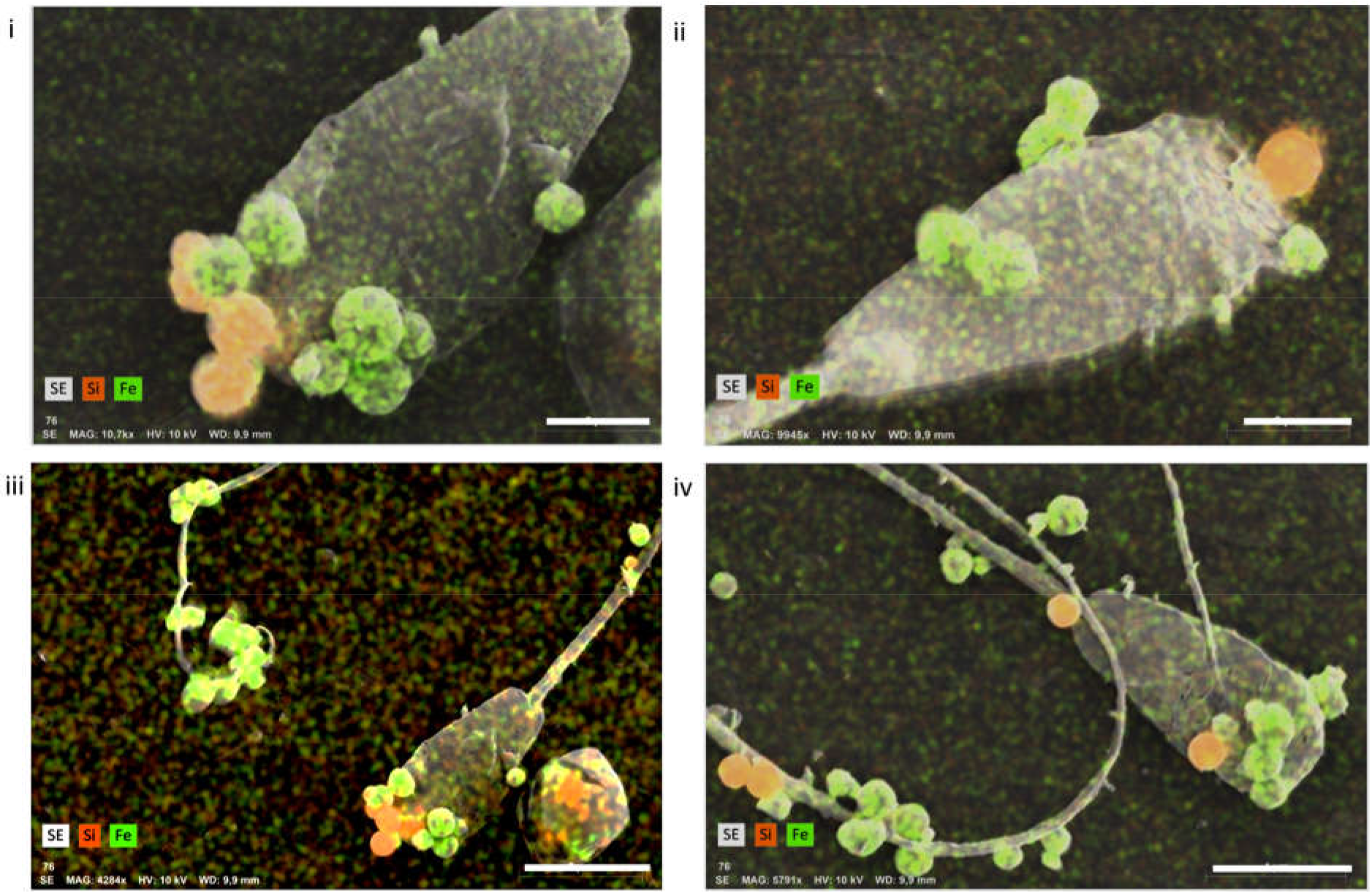
Superimposed cryo-SEM (SE) and EDX line scans confirming the presence of silica (Si) and iron oxide (Fe) particles on bovine sperm. Scale bars are in i) and ii) 2 μm, in iii) and iv) 5 μm.

In conclusion, we demonstrate an approach to use charge-based interactions between microparticles and sperm cells to map the charge distribution of the cell’s membrane. This is a straightforward procedure because it does not involve advanced surface modification techniques. Although the net surface zeta potential is commonly being used to separate sperm cells,^[13]^ the distribution of charge across the sperm cell’s membrane has been overlooked so far. It is worth investigating to understand the dynamic change of the sperm’s surface, all the way to the event of fusing with the oocyte. During sperm maturation, the sperm membrane organization changes dramatically which results in the capacitated state, which, in turn, allows the sperm to bind to the zona pellucida and acrosome react.^[24]^ The fusion between sperm and egg is known to be a process involving abrupt changes of electrical charge that allow the electrical blockage of the oocyte by reversion of its surface charge to avoid polyspermy.^[17,18]^ Our findings of negatively charged particles binding especially to the sperm heads suggest that positively charged regions exist on the sperm’s membrane. These positively charged regions disappear in maturated sperm, observed in this work by loss of binding to negatively charged particles. This loss of positive surface charge might derive from the fact that capacitation involves a decrease of the NNC, removal of cholesterol from the phospholipid bilayer and increase of membrane permeability.^[25]^

Pursuing these proof-of-concept experiments further in the future by attaching smaller particles to the sperm’s surface will give a more detailed charge map, but the easy read out by optical microscopy will have to be replaced by electron microscopy. It will also be interesting to compare sperm cells of different developmental stages (e.g., epididymal sperm vs. ejaculated, pre- and postacrosomal sperm) to investigate the dynamic changes of the sperm and when and why the sperm’s surface charge reverses from negative to positive.^[13]^ Since the interaction between surrounding media and cells also contribute to the change of membrane composition of sperm, it will be crucial to look at different media compositions and their effect on the sperm’s surface charge distribution. These complex endeavors will be significantly enriched by computational simulations that allow the modelling and thereby prediction of sperm-particle interactions.

## Experimental Section

### Sperm preparation

Cryo-preserved bovine semen was purchased from Masterrind GmbH and stored in liquid nitrogen until use. Bovine semen straws were thawed in a 38°C water bath for 2 minutes, resuspended in 1mL SP-TALP (Caisson Lab, modified tyrode’s albumin lactate pyruvate medium) and centrifuged at 300g for 5 minutes in SOFT mode. The sperm sample was then resuspended in 0.5mL fresh SP-TALP for viability studies or in water for particle-binding studies.

### Preparation of particles

*H1:* 90mL of 6M NaOH solution was added quickly to 100ml of 2M FeCl_3_*6H_2_O solution, the solution was stirred for 10min and 10ml of MilliQ water were added. The resulting mixture was added into a pyrex bottle and kept for 10 days at 100°C. The product was washed several times with ethanol and MilliQ water and subsequently characterized.

*H2:* To obtain nanoscale iron oxide particles a synthetic strategy from Baeza et al.^[26]^ was followed. Briefly 10.21g of FeCl_2_*4H_2_O (0.052mol) were dissolved in 1L of MilliQ H_2_O, pouring a second solution of FeCl_3_*6H_2_O (28.35g, 0.104mol) in 57mL of HCl 1.5mol*L^−1^ under strong stirring, yielding nanometric magnetite (Fe_3_O_4_). The black flocculate was dispersed to a 2mol*L^−1^ HNO_3_ solution and stirred for 2-3min. After decantation, the particles were oxidized in a solution of Fe(NO_3_)_3_*9H_2_O in H_2_O (0.3366 mol*L^−1^) at 100°C, keeping that temperature for 30min. Another decantation (2-3h) was carried out and the product was dispersed in a 0.02mol*L^−1^ HNO_3_ solution and used without further purification.

*H3:* Hematite particles were synthetized by a modified recipe from Demarchis et al.^[27]^. In brief, a 5.4M NaOH solution was mixed with a 2M FeCl_3_*6H_2_O solution, stirred and after addition of 2mL of a 2M solution of nitriloacetic acid the solution was aged at 100°C for 7 days. Subsequently, the particles were washed with water and ethanol and redispersed in MilliQ water.

*H4:* Rice-shaped particles were synthesized following a procedure by Ohmori et al.^[28]^. A solution of FeCl_3_*6H_2_O (0.02 mol*L^−1^) was aged with a NaH_2_PO_4_ solution (0.0004 mol*L^−1^) during 72 hours in a Pyrex bottle at 100°C. The obtained product was centrifuged and washed with acetone.

*Silica coverage* was previously described by Rufier et al.^[29]^.

*SiO_2_@H1:* 20mg H1 particles were dispersed in a PVP solution in H_2_O (400mg*44mL^−1^) and sonicated during 2h followed by 15h of agitation. Particles were centrifuged and washed with water to remove excess PVP. After redispersion in 2.5mL H_2_O particles were added to 45mL EtOH, sonicated, and stirred at 400rpm. After 30min TMAH in EtOH (0.1865mol*L^−1^) was added, followed by 30min of stirring. 50μL of APTES were diluted in 200μL of EtOH and added dropwise. After 30min of stirring a mixture of APTES, TEOS and EtOH (4:150:1000) was added in 3 portions each 385μL.

*SiO_2_@H4:* 20mg H4 particles were treated as described for SiO_2_@H1.

*S1:* Silica spheres were produced according to a previously published two step method.^[30]^ To obtain seeds 1.9mL TEOS were diluted in 13mL propanol. 0.7mL NH_3_ (25%) diluted in 1.4mL*H_2_O were added at room temperature and subsequently aged for 60min at 50°C.

To this solution 7mL TEOS diluted in 300mL propanol were added under constant stirring. Before heating to 50°C 20mL*NH_3_ (25%) in 40mL of water were added under constant stirring. As last step 120mL of TEOS were added at an addition rate of 0.5g*min^−1^ and then allowed to age for 1h. The product was washed with water and ethanol and dried prior to usage.

The Zetapotential of all particles was determined with the Zetasizer Nano ZSP (Malvern Panalytical GmbH, Kassel, Germany) and are displayed in Table 1.

### Sperm-particle-binding assays

Particle suspensions were vortexed and well suspended by sonicating for 10min before adding 50μL of this particle suspension to 50μL of sperm suspension. The mixture was incubated at 38°C for 30min before analysis. A drop of 10μL of the sperm-particle suspension was transferred to a glass slide, covered with a cover slide and immediately observed under an inverted microscope Leica DMi8 at 20x, 40x or 100x magnifications (Leica Microsystems GmbH, Wetzlar, Germany). A total of 100 cells was analyzed for each particle type and 3-5 repetitions were done for each type of particle.

### Sperm maturation

Cryo-preserved bovine semen was thawed as mentioned above and centrifuged in SP-TL (Caisson Lab, no pyruvate or albumine) at 300g for 5 minutes and resuspended in 250μL SP-TALP supplemented with 100μg*mL^−1^ heparin-fluorescein conjugate H7482 (Thermo Fisher Scientific, Fisher Scientific GmbH, Schwerte, Germany) and incubated for 30min at 37°C prior to particle binding assay.

### Zeta potential measurements of bovine sperm cells

Custom-made microelectrophoresis was used to measure the bull sperm charge by applying an electric field to the sperm cell suspension in water. The electrophoresis cell consists of a U-shape glass chamber filled with a sperm medium. Platinum electrodes were used to apply a current, generated from the power supply, to the electrophoresis cell. Six milliliters of sperm medium were added to the electrophoresis chamber with 50μL of sperm cells for each measurement. Voltage of 20 V was applied and from the trajectories of the sperm cells, the electrophoretic mobility was calculated from the sperm velocity due to electric field divided by applied voltage multiplied per unit length of microelectrophoresis chamber. Subsequently, the Zeta potential (ζ) could be calculated from the equation.

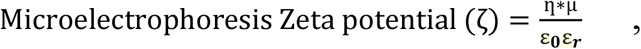

where η is water viscosity, μ is the electrophoretic mobility, ε_0_ is vacuum permittivity and ε_r_ is the dielectric constant. Tracking of 100 sperm cells at 20 V gave an average zeta potential of –13mV.

Surface zeta potential of bovine sperm cells was also estimated by zetasizer Nano ZSP. A dilution of sperm medium (1:100) (TL sperm media (no albumin): deionized water) with washed sperm cells was filled into a 1mL cuvette and inserted into the zetasizer for measurement (see Figure S1 for exemplary result). Six measurement runs were conducted and resulted in average zeta potential of −27.4 mV for bovine sperm cells in 1:100 diluted medium without albumin.

### Sperm viability test

The Live/Dead Sperm Viability Kit was purchased from Molecular Probes L7011 (Thermo Fisher Scientific, Schwerte, Germany). SYBR 14 (green dye, stains live cells) was diluted 1:50 in SP-TALP. 1μL of this dilution was added to 100μL of sperm-particle suspension. Subsequently, 0.5μL propidium iodide (red dye, stains dead cells) was added to the sperm-particle suspension. The sample was incubated at 38°C in darkness for at least 5min before fluorescent imaging.

### Cryo-SEM & EDX

For cryo-SEM of sperm-particle suspensions, bovine sperm cells and particles were washed separately twice in deionized water before combining them for attachment assays. They were incubated for 30min at a ratio of 1:1:1 (e.g., 3μL iron oxide particles:3μL silica particles:3μL sperm pellet) in 100μL deionized water. The cryo-(FE-)SEM SUPRA 40VP-31-79 (Carl Zeiss SMT Ltd., Oberkochen, Germany) equipped with an EMITECH K250X cryo-preparation unit (Quorum Technologies Ltd., Ashford, Kent, United Kingdom) was used for imaging 1μL of the sperm-particle suspension, which was dropped onto glass or aluminum tape (in case of EDX) fixed to the cryo-SEM holder and immediately shock-frozen in liquid nitrogen in the slushing chamber. From there it was transferred to the cryo preparation chamber at −140°C, sublimed for 10-15min at −70°C, and sputter coated with platinum (layer thickness ca. 6nm). Subsequently, it was transferred to the SEM, and then examined in a frozen state at 5kV accelerating voltage and −100°C temperature using the secondary electron (SE) detector. In addition, the EDX detector Bruker x-flash 6/100 and Esprit 2.0 Software (Bruker Corporation, Billerica, MA, USA), connected to the cryo-(FE-)SEM, were applied to carry out energy-dispersive x-ray analyses at a working distance of 7-10mm, at 10 and 20kV acceleration voltage, 51200 income counts per second (cps), and a measurement period of 60s, detecting a total (element) percent by weight of 83 and a clear presence of Si and Fe (Fig. 7). The aluminium tape substrates were used, because the EDX spectra of these materials did not interfer with the characteristic X-ray peaks of the targeted elements silicon and iron oxid.

## Acknowledgements

The authors thank the Chair of Applied Zoology and the Chair of Botany at the faculty of Biology of TU Dresden, Germany, for providing lab space & equipment and access to cryo-SEM/EDX, respectively. VM and JG thank the Zukunftskonzept of the TU Dresden, funded under the Excellence Initiative of the German Science Foundation (DFG) for funding. JS and PS thank the Volkwagen foundation for a Freigeist fellowship (grant number 91619) and the SMWK for the BacMot support.

---

**In this article, the attachment of differently charged microparticles to bovine sperm cells is investigated**. The binding of positively and negatively charged particles to different distinct regions of the sperm’s surface is promising for charge mapping, which could give insight into the dynamic change of surface charge, an important indicator for changes during a sperm’s development.

**Sperm, microparticle, membrane, charge**

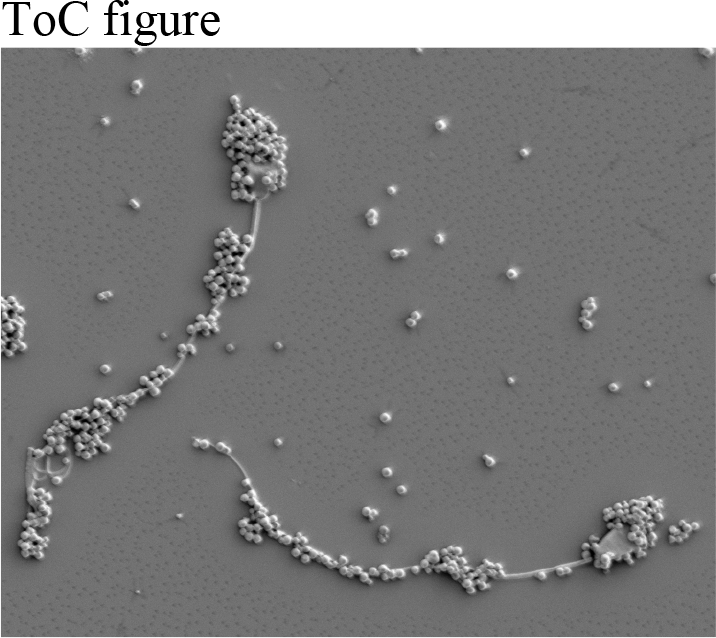

## Supporting Information

**Figure S1:**
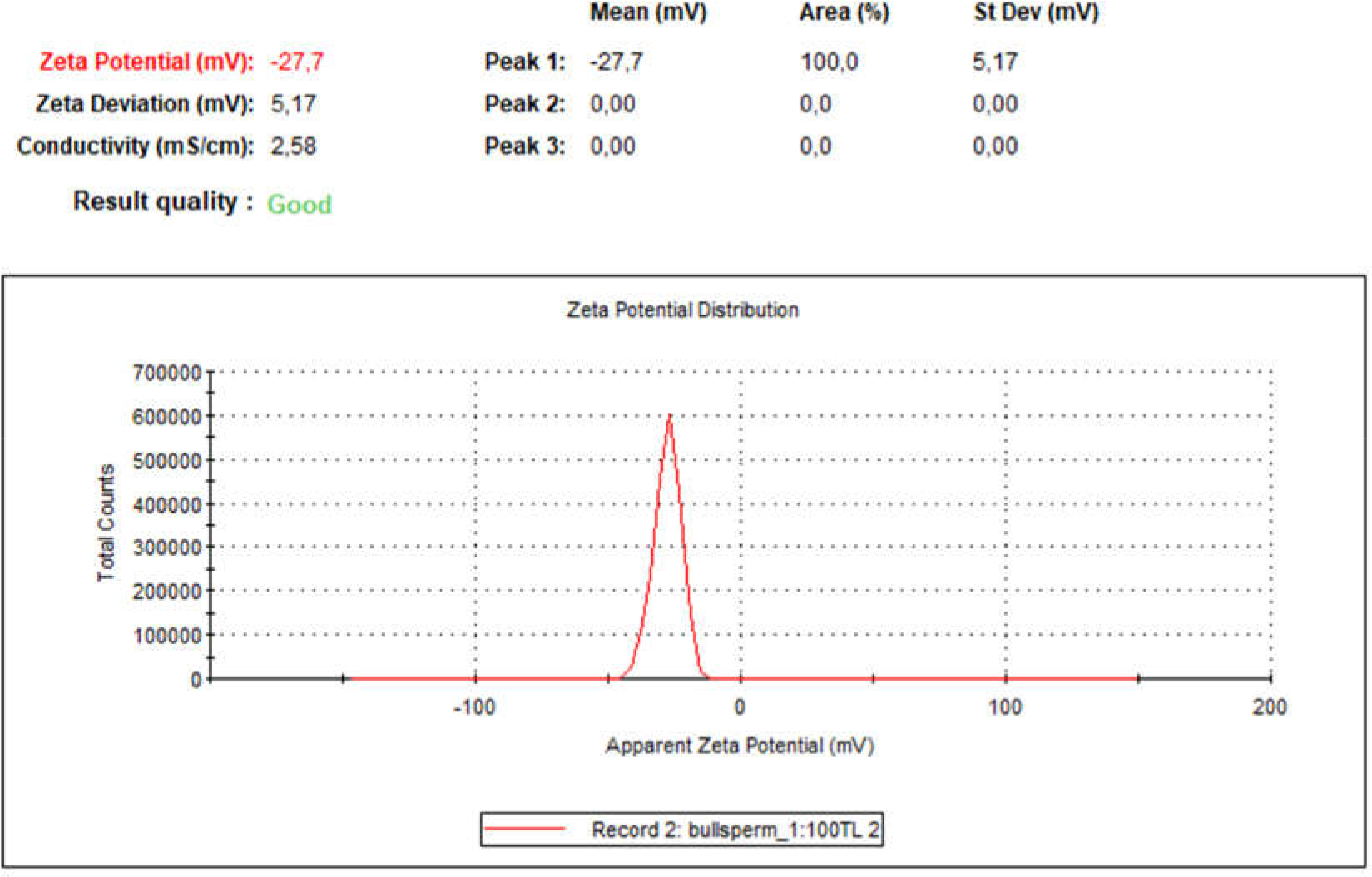
Example of Zeta potential measurement of bovine sperm cells by the Zetasizer Nano ZSP (Malvern Panalytical GmbH, Kassel, Germany).

